# Accelerated amyloid deposition in SARS-CoV-2 infected mouse models of Alzheimer’s disease

**DOI:** 10.1101/2024.12.09.627570

**Authors:** Parag Parekh, Andrew A. Badachhape, JeAnna R. Redd, Lauren J. Bonilla, Prajwal Bhandari, Alexander R. Kneubehl, Rohan Bhavane, Jennifer L.S. Clinton, Prasad Admane, Renuka Menon, Mayank Srivastava, Xianwei Sun, Saphal Narang, Eric Tanifum, Ketan B. Ghaghada, Shannon E. Ronca, Ananth V. Annapragada

## Abstract

Familial Alzheimer’s disease (AD) involving known AD causing genes accounts for a small fraction of cases, the vast majority are sporadic. Neuroinflammation, secondary to viral infection, has been suggested as an initiating or accelerating factor. In this work we tested the hypothesis that SARS-CoV-2 (SCV2) viral infection accelerates the development of AD pathology in mouse models of AD. We profiled transcriptomic changes using transgenic APP/PSEN1 and P301S mouse models that develop AD pathology and k18hACE2 mice that express the humanized ACE2 receptor used by SCV2 to enter cells. This study identified the interferon and chemokine responses constituting key shared pathways between SCV2 infection and the development of AD pathology. Two transgenic mouse models of AD: APP/PSEN1 (develops amyloid pathology) and 3xTg AD (develops both amyloid and tau pathology) were crossed with k18-hACE2 mice to generate hybrid hACE2-3xTg and hACE2-APP/PSEN1 mice. Neuroinflammation and amyloid deposition in the brain of infected mice were imaged *in vivo* using molecular MRI (mMRI) probes and confirmed postmortem by histopathology. Results show that 11-14-month-old SCV2 infected hACE2-3xTg mice exhibit neuroinflammation 10 days post infection and 4–5-month-old hACE2-APP/PS1 hybrid mice develop amyloid deposits, while age-matched uninfected mice exhibit neither phenotype. This suggests that SCV2 infection could induce or accelerate AD when risk factors are present.

## Introduction

COVID-19 and Alzheimer’s disease (AD) both disproportionately affect the elderly (age ≥ 65) and were the third and seventh leading causes of death in the USA in 2021 as reported by the CDC^1^. The vast majority of AD cases are sporadic, with no known genetic underpinning^2,3^. Neuroinflammation secondary to viral infection has long been suspected as a contributing factor driving the development of AD^4–8^.

Increasing evidence links the overlapping molecular pathways of COVID-19 and neurodegenerative disease including AD. A recent cross-sectional study^9^ studied 310 COVID-19 patients and controls with other CNS inflammatory disease and demonstrated Blood Brain Barrier (BBB) impairment, elevated microglial activation, and decreased regional brain volumes in COVID-19 patients. A transcriptomic study of 22 patients with COVID-19 and other viral diseases showed dysregulation of both choroid plexus cell types and upper layer excitatory neurons^10^ in COVID-19 patients. Similarly, microbleeds^11^ and microvascular injury^12^ have been reported in COVID-19 patients. A causal effect of COVID-19 on Alzheimer’s disease (AD) has been postulated from a Mendelian randomization study^13^ of a large European cohort. Similarly, a TriNetX based study of 6 million adults in the US concluded that COVID-19 patients (age >65 years) had a significantly higher probability of receiving a new diagnosis of AD in the 12 months following infection. Elevated levels of neurofilament light (NfL), and decreased CSF soluble-APP, soluble-APPβ, Aβ40, Aβ42 and Aβ40/Aβ42 ratio were observed in COVID- 19 patients compared to uninfected controls^14^. Signaling pathways in common with AD were observed in COVID-19 patients by a transcriptomic study^15^. A spatial transcriptomic study coupled with post mortem histopathology of COVID-19 patients^16^ suggested significant overlap of activated pathways with both Alzheimer’s and Parkinson’s diseases, and showed elevated amyloid deposition but not tau pathology. Common neurological and ophthalmological manifestations of AD and COVID-19 have been described^17,18,19^. Separately, other viruses have also been implicated, foremost is HSV-1, linked to the initiation of an AD phenotype^20–24^.

In this work, we investigate neuroinflammation caused by SCV2 infection and its role in neurodegeneration in mouse models of AD. The phenotype of infected mice was used as a model to study infection induced pathogenesis in the context of predisposition to the disease. SCV2 infects humans via the ACE2 receptor but does not bind mouse ACE2. A humanized ACE2 transgenic mouse was previously generated(k18-hACE2)^25^ to provide a mouse model of productive infection. SCV2 infected k18-hACE2 transgenic mice develop lung inflammation, cytokine-storm and neuro-invasion similar to human COVID-19^26,27^. To test the effect of infection on AD, we first generated two SCV2 susceptible AD mouse models by cross breeding *k18-hACE2* mice with established AD models: (1) APP/PSEN1^28^, a transgenic mouse model that expresses mutant human amyloid precursor protein and a mutant human presenilin-1 associated with early onset AD that develops amyloid plaque pathology and seizures; (2) 3xTg-AD^29^ mice bearing both mutant APP and tau (P301L) developing age-dependent tau and amyloid pathology and a cognitive deficit phenotype preferentially in female mice. Genotypes of the cross- bred mice were verified for the individual transgenes. We conducted a comprehensive profiling of the brains of APP/PSEN1, P301S, and k18-hACE2 models using bulk RNA sequencing (RNA-Seq) to investigate the overlap in transcriptomic alterations associated with ancestral SCV2 infections and AD. Mice were then aged to a point where frank AD pathology (amyloid in APP/PSEN1 mice and both amyloid and tau in 3xTg mice) were not expected, and infected with the Delta variant of SCV2 as a model of severe COVID-19^30^. Pathological amyloid, tau and neuroinflammation were studied in hACE2-APP/PSEN1 and hACE2-3xTg AD mice by both molecular MRI (mMRI) and histopathology.

## Methods

### Animal Models

All animal studies were approved by the Institutional Animal Care and Use Committee. Individual breeding pairs of APP/PSEN1(034832-JAX), 3xTg-AD (034830- JAX) and k18-hACE2 (034860-JAX) mice were purchased from the Jackson Laboratory (JAX). Hemizygous transgenic male APP/PSEN1 and homozygous 3xTg-AD mice were crossed with female k18-hACE2 mice. Offspring genotypes for these include the quadruple-transgenic hACE2-3xTg, triple-transgenic hACE2-APP/PSEN1 and the hemizygous and non-transgenic hACE2^WT^-3xTg, hACE2^WT^-APP/PSEN1^Tg^, hACE2^Tg^- APP/PSEN1^WT^, hACE2^WT^-APP/PSEN1^WT^. Pups were weaned at 21 days. Ear punch tissue was used for genotyping by PCR using primers for the transgenes hACE2, MAPT, APP and PSEN1 as recommended by JAX, with amplified products visualized on 1.5% agarose gel.

Female quadruple transgenic hACE2-3xTg offspring identified by genotyping were allowed to age for 11-14 months before infection and mMRI. Similarly, triple transgenic hACE2-APP/PSEN1 mice, both male and female, were aged to 3-4 months before infection and mMRI studies. These ages were chosen to be younger than those at which frank AD pathology (amyloid in the case of APP/PSEN1; both amyloid and tau in the case of 3xTg) have been historically observed^28,29^.

All animals were allowed free access to water and food and provided with a 12 h light/dark cycle.

### SCV2 Infection-delta variant

hACE2-3xTg (n=16, 11-14months) and hACE2-APP/PSEN1 mice (n=27, 3 months) were moved from the main vivarium (ABSL2) into ABSL3 containment and acclimated to the environment for 3 days. SCV2 delta variant was obtained through BEI Resources, NIAID, NIH: SARS-Related Coronavirus 2, Isolate hCoV-19/USA/PHC658/2021 (Lineage B.1.617.2; Delta Variant), NR-55611, contributed by Dr. Richard Webby and Dr. Anami Patel. Mice were then intranasally infected with 1x10^3^ PFU of the SCV2 Delta variant b.1.617.2. Body conditioning scores and weights were recorded daily for all mice during the experimental period (Supplementary Figures 1-2). Humane euthanasia was carried out when mice scored 4 twice in one 24-hour period or scored 5. Short term, acute study in hACE2-3xTg mice were performed as follows: on day 5 post infection, mice underwent a non-contrast baseline set of MRI scans and were injected with liposomal gadolinium contrast on day 6. Delayed post-contrast images were collected on day 10. Long term study in hACE2-APP/PSEN1 mice was carried out as follows: 30 days post infection, mice underwent a non-contrast baseline set of MRI scans and were injected with liposomal gadolinium contrast on day 26. Delayed post-contrast images were collected on day 30. Uninfected hACE2-3xTg mice and hACE2-APP/PSEN1 mice (6 each) were housed in the main vivarium and imaged using identical pre-contrast and post-contrast protocols and sequences.

### Transcriptomics

Six- to eight-week-old k18-hACE2 mice (n=8) were intranasally infected with 1x10^5^ plaque forming units (PFU) to induce severe disease or mock infected with sterile PBS (n=6). Mice were euthanized 4-11 days post infection (dpi). Brains were harvested and snap frozen. Brain tissue was homogenized using a bead mill tissue homogenizer TissueLyser II (Qiagen, CA). Brain homogenate was inactivated in Trizol prior to removal from the BSL-3 facility. RNeasy Lipid Tissue kit (Qiagen, CA) was used with on-column DNase digestion as detailed in the kit user guide. Resultant RNA was measured by NanoDrop (ThermoFisher, MA). Poly(A) RNA sequencing library was prepared following Illumina’s TruSeq-stranded-mRNA sample preparation protocol. RNA integrity was checked with Agilent Technologies 2100 Bioanalyzer. Paired-ended sequencing was performed on Illumina’s NovaSeq 6000 sequencing system by LC Sciences, Houston, TX. Reads with adaptor contamination, low quality, and undetermined bases were removed using Cutadapt^31^ and in-house perl scripts. Sequence quality was verified with FastQC. HISAT2^32^ mapped reads to the mouse genome. StringTie^33^ assembled the mapped reads, and all transcriptomes were merged using perl scripts and gffcompare. Expression levels of transcripts were estimated with StringTie and ballgown(http://Bioconductor.org). StringTie calculated mRNA expression levels (FPKM). Differential expression analysis was performed using DESeq2^34^ for groups and edgeR^35^ for samples. mRNAs with FDR < 0.05 and absolute fold change ≥ 2 were considered differentially expressed. Ontological and pathway analysis was performed using Enrichr^36^ and functional term enrichment was performed using RummaGEO^37^ with genes with nominal p-value less than 0.05 to increase analytical power. All visualizations of RNA-seq, differential expression analysis, and ontological analysis data were generated using Microsoft Excel.

### Preparation of molecular MRI probes

Liposomal nanoprobes targeted to the folate receptor and to amyloid plaques, and non-targeted particles to image vascular leak and to serve as controls were prepared by hydration-extrusion, using methods described in detail previously^38,39^. In brief, lipids (HSPC, Cholesterol, DSPE-DOTA-Gd, mPEG2000-DSPE) were first dissolved in ethanol. For targeted particles (folate targeted, amyloid targeted), an additional DSPE-PEG3400- (targeting ligand) compound was added. A lipid fluorescent dye, DiI, was added to the lipid mix for post-mortem fluorescence microscopy analysis. The ethanolic solution of all lipids was hydrated with a saline buffer (ethanol:saline volume ratio = 0.1) and then extruded through nucleopore membranes to produce liposomes with diameter ∼150nm as measured by dynamic light scattering. Inductively coupled plasma optical emission spectroscopy (ICP-OES) was performed to measure gadolinium and phosphorus (phospholipid) content of liposomal formulations. Liposomal composition and ligand structures are shown in Figure 4. For amyloid targeted particle studies without the DiI dye, we utilized ADx-001, which was otherwise identical in composition, and was provided by Alzeca Inc.

### Magnetic Resonance Imaging (MRI)

Mouse imaging in the biosafety facility (ABSL3) was performed on a 1T permanent MRI scanner (M2 system, Aspect Imaging, Shoham, Israel) while non-infected imaging occurred outside the ABSL3 on a separate 1T permanent magnet MRI scanner (M7 system, Aspect Imaging, Shoham, Israel). Both scanners used a mouse head coil for neuroimaging. All mice underwent the same T1-weighted spin echo sequence with the following parameters: TR: 600 ms, TE: 11.5 ms, slice thickness: 1.2 mm, matrix: 192x192, FOV: 30 mm, slices: 16. During imaging, mice were sedated using 3% isoflurane, placed on the MRI animal bed, and then maintained at 1-2% isoflurane delivered using a nose cone setup. Breathing rate was monitored through a pressure pad placed below the abdomen.

3xTg mice and hACE2-3xTg mice were imaged 10 days after infection, using the folate-targeted nanoparticle agent (see Figure 3) as a probe of neuroinflammation. For this study, 5 days after infection (or sham instillation for uninfected controls), a pre-contrast T1-weighted MR scan was obtained. On day 6, folate targeted nanoparticle contrast agent (0.2 mmol Gd/kg) was injected i.v. via tail vein for detection of neuroinflammation. On day 10, the mice were once again imaged using the same T1- weighted sequence. Non-targeted (no folate ligand) control nanoparticles were injected into control group mice which were in turn imaged in identical fashion.

APP/PSEN1 and hACE2-APP/PSEN1 mice were imaged 30 days after infection, using an amyloid targeted nanoparticle agent (AβNP) (see Figure 3) as a probe of amyloid deposition. 25 days after infection (or sham instillation for uninfected controls), a pre-contrast T1-weighted MR scan was obtained. On day 26 the amyloid targeted nanoparticle contrast agent (AβNP) (0.2 mmol Gd/kg) was injected i.v. via tail vein for detection of amyloid pathology. On day 30, the mice were once again imaged using the same T1-weighted sequence. Non-targeted (no amyloid-targeting ligand) control nanoparticles were injected into control group mice which were in turn imaged in identical fashion.

### Histopathology

Uninfected mice, after euthanasia, were perfused with 0.9% saline followed by 4% paraformaldehyde and harvested brains were then immersion-fixed in 4% formaldehyde for 48 h at 4 ⁰C, transferred to 30% sucrose for cryoprotection and embedded in OCT. Brains of SCV2 infected mice were immersed in 10% neutral buffered formalin with formalin change occurring every 24 hours three times to verifying viral inactivation. Downstream processing was consistent with uninfected mice. One hemisphere was used for histopathology and the other was used to generate brain homogenates for bulk RNA sequencing studies. Brain sections were cut at thickness of 50 µm. Microwave (GE, 1200W) assisted antigen retrieval in citrate buffer (pH=8.5) was performed in a microwaveable tray (RPI, Mt. Prospect, IL #248270) 2X for 5 minutes at power level 4 with one minute interval, followed by 15 minutes cooling time. The sections were stained overnight at 4⁰C with 1:100 dilutions of the following antibodies: p-tau antibody (Thermofisher, AT8) recognizing p-tau Ser202/Thr205, amyloid antibody (Biolegend, β amyloid 4G8) recognizing amino acid residues 17-24 of β amyloid and IBA1 (Wako Fujifilm, IBA1) specific to activated microglia. Sections were washed with PBS 3X and then incubated for 2hrs with a 1:1000 dilution of appropriate secondary antibody. DAPI staining was performed after PBS washing. Fluoromount G (Electron Microscopy Sciences, 17984) was used to mount slides which were visualized on an Olympus Fluoview LV100 confocal microscope at 40x or 60x magnification.

### Data and statistical analysis

All MR images were analyzed using a combination of thresholding and segmentation in OsiriX (version 5.8.5, 64-bit) and MATLAB (version 2015a). The signal change between pre-contrast and delayed post-contrast images was evaluated by quantifying signal intensity (SI) in cortical regions near the center of the image stack^40^. All statistical analysis was performed using Students t-test.

List of Antibodies - AT8 (Thermo Fisher Scientific, Waltham, MA, #MN1020), β amyloid (BioLegend, San Diego, CA, #800711), IBA1 (Wako FujifIlm, Richmond, VA, #019- 19741), AF488 (Invitrogen, Waltham, MA, #A11008), AF647 (Invitrogen, Waltham, MA, #A32728TR)

## Results

### Comparison of transcriptomic profiles following ancestral SCV2 infection and in AD mouse models

We conducted transcriptomic profiling using bulk RNA-Seq on brain homogenates of 6–8-week-old k18-hACE2 mice that were infected with a lethal dose of ancestral SCV2 and euthanized at 4-11dpi alongside mock-infected age-matched controls. Profiling of AD mouse models was carried out on 8-month-old APP/PSEN1 and 12-month-old P301S transgenic mice and wild type sibling controls. The profiling yielded 1,706 (190 up and 1,516 down) differentially expressed genes between the SCV2 infected and uninfected group, 265 (134 up and 131 down) for the APP/PSEN1 and 2,150 (1,243 up and 907 down) for the P301S transgenic over wild-type groups (p<0.05). Differential gene expression is plotted as heat-maps in Figure 1A showing the intersection of genes that are common between ancestral SCV2 infection and AD development. Figure 1B shows the activated pathways for the differentially expressed genes in-common using BioPlanet2019, mSigDB Hallmark 2020 and Reactome 2022 Mouse suggesting that the inflammatory and chemokine responses are the processes in common between SCV2 infection and AD pathology development in these mouse models. Similarly, the gene ontologies for biological processes and molecular function are represented as a TreeMap in Figure 1D suggest immune and chemokine responses among the shared differentially expressed genes. A functional enrichment of the terms across published enriched mouse gene sets (Figure 1C) was also consistent with the inflammatory processes being a shared characteristic of ancestral SCV2 infection and AD.

**Figure 1:**
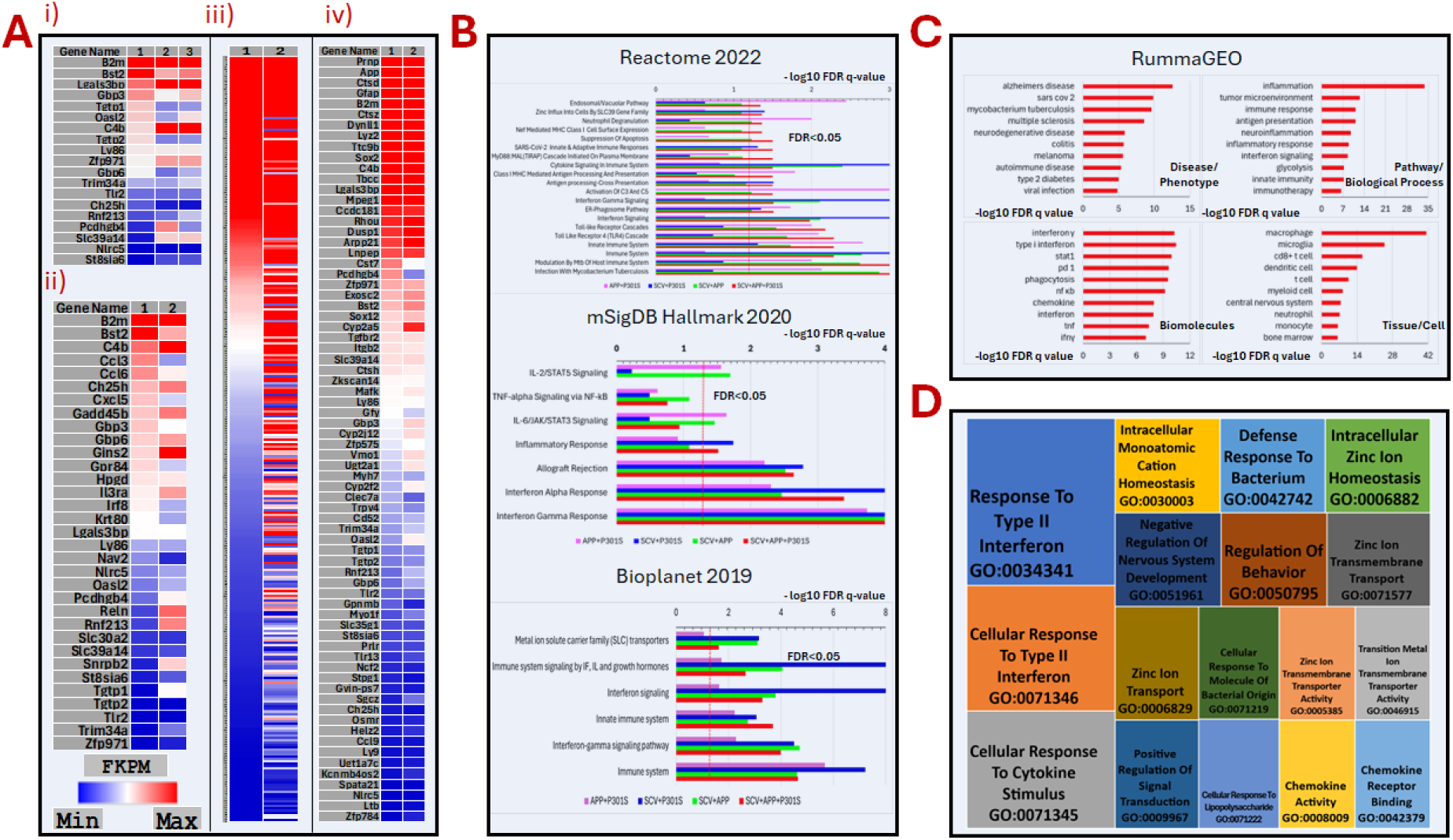
Bulk RNAseq of whole mouse brain homogenates yields insights into pathways in common between a mouse model of tau deposition (P301S), amyloid deposition (APP/PSEN1) and ancestral SCV2 infection at 4 days post infection in k18-hACE2 mice. A. Heat map shows the intersection of differentially expressed gene sets (corrected p<0.05) between infected and uninfected mice, and between P301S and APP/PSEN1 transgenic and wild type mice. (i) SCV2 (1), APP/PSEN1(2) and P301S(3); (ii) SCV2 (1) and APP/PSEN1(2); (iii) SCV2(1) and P301S(2); (iv) APP/PSEN1 and P301S . B. Differentially expressed genes in-common demonstrate activated pathways (-log10q- value>1.3) in the Reactome 2022, mSigDB Hallmark 2020 and BioPlanet2019. Of note, inflammatory processes represented by interferon signaling and the complement cascade constitute many of the common pathways. Also of note, the APP/PSEN1 mice demonstrate Type 1 interferon and cell-specific innate immune response that are characteristic of SCV2 infection. Taken together, these results suggest inflammatory/cytokine processes in common between SCV2 infection and Alzheimer’s disease in these mouse models. C. The enriched genes from SCV2+APP/PSEN1+P301S were compared to existing mouse gene sets using RummaGEO identifying common terms in matching gene sets. Functional term significance was identified for the Disease/Phenotype, Biomolecules, Tissue/Cell and Pathway/Biological Process and confirmed the activation of inflammatory processes characteristic of infection and Alzheimer’s disease. D. Tree Map visualizing the significant gene ontologies (FDR q-val <0.05) GO: Biological Process and Molecular Function are heavily concentrated in the immune and chemokine response branches. Other highly represented branch is Zinc ion homeostasis and transport activity.

### Characterization and infection of hybrid hACE2-APP/PSEN1 and hACE2-3xTg AD mice

The progeny of k18-hACE2 X 3xTg AD and k18-hACE2 X APP/PSEN1 matings (Figure 2A) were genotyped and compared with the parental strains to confirm successful generation of the hybrid hACE2-3xTg-AD and hACE2-APP/PS1 mice. Figure 2Ai illustrates the outcomes of the crossings and 2Aii shows the agarose gel images for transgenes present in representative parents and hybrid progeny and confirms presence of all relevant transgenes in the crossbred mice.

**Figure 2.**
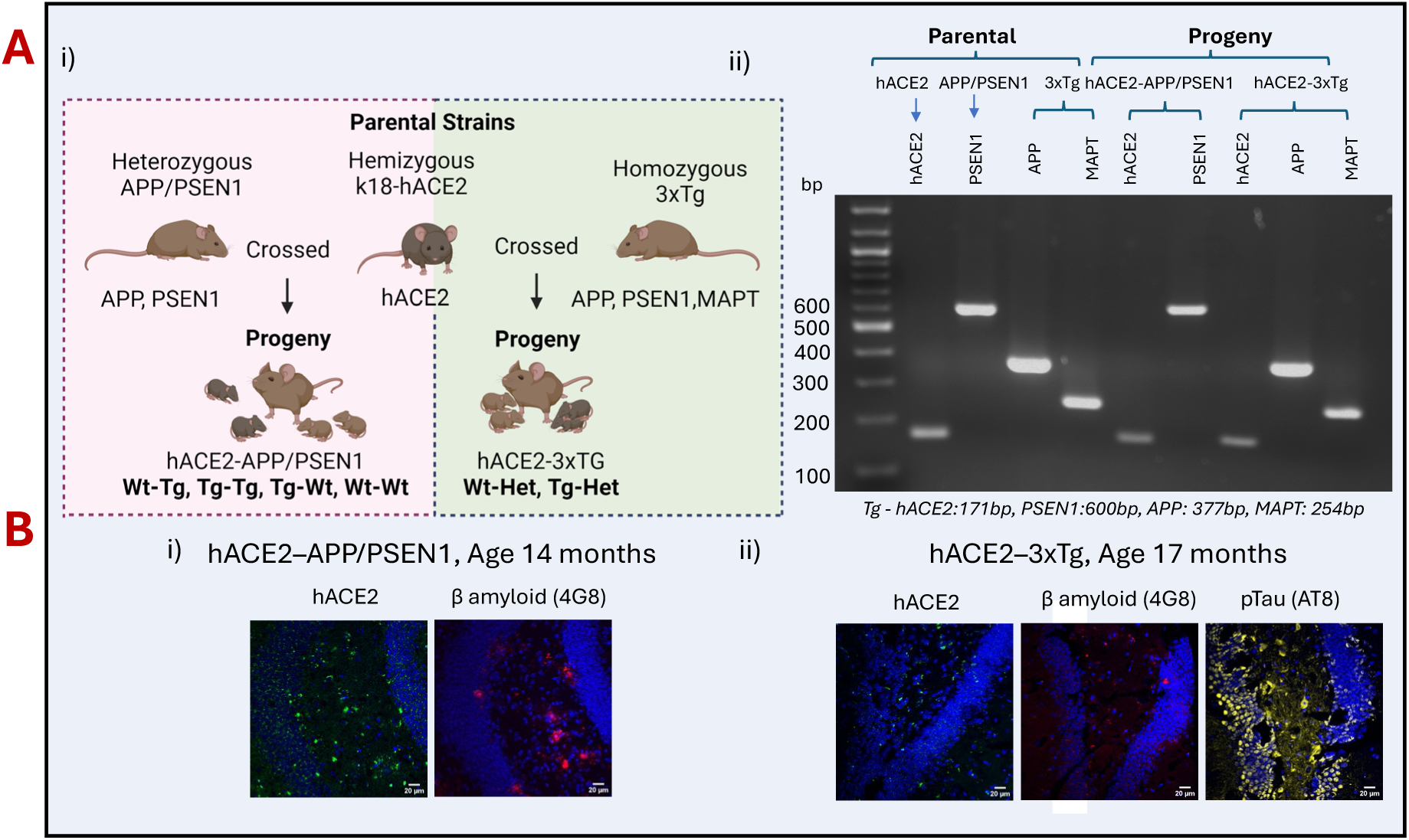
Generation of a mouse model that can be infected with SCV2 and shows AD phenotype. (A) K18-hACE2 hemizygous mice were bred with hemizygous APP/PSEN1 and homozygous 3xTg-AD mice to yield progeny the hybrid transgenic mice, hACE2- APP/PS1 with transgenes ACE2, APP and PSEN1 and the quadruple transgenic mice hACE2-3xTg with transgenes ACE2, APP, PSEN1 and MAPT. (ii) Genotyping the parental and hybrid mice for the presence of transgenes in both the hybrid mouse models. (B) Histopathological characterization of the crossed strains confirming the presence of AD related pathology. Mice ages were selected based on the reported confirmed presence of phenotype in the relevant parental strains. i) Hybrid hACE2-APP/PS1 mice at age 14months (three transgenes) and ii) Hybrid hACE2-3xTg mice (four transgenes) at age 17months, stained for transgenic hACE2 with antibody (green) and amyloid plaques stained by the amyloid 4G8 antibody (red) and phosphorylated Tau stained by antibody AT8 (gray) in the brains of representative animals.

**Figure 3:**
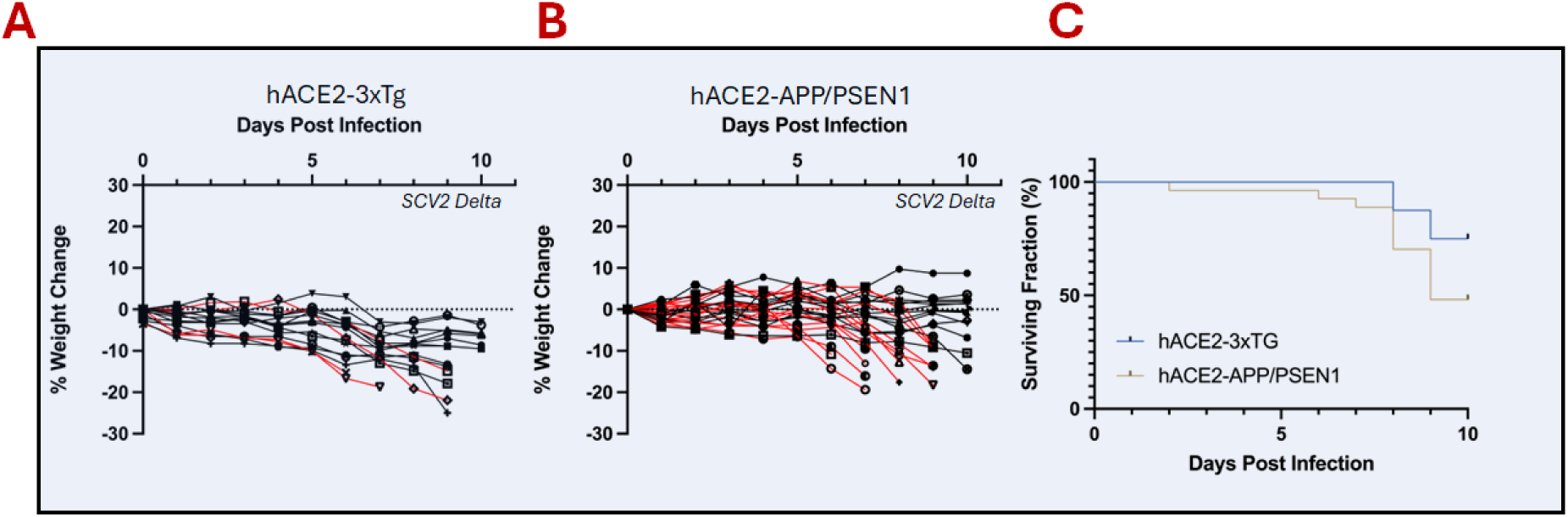
Susceptibility of hybrid hACE2-3xTg and hACE2-APP/PS1 mice to SCV2 Infection. Both the hybrid mouse models were infected with SCV2 delta variant b.1.617.2 at 1x103 PFU. Body weight changes at 10days post infection are plotted for the (A) hACE2-3xTg mice with 4 of 16 mice succumbing to SCV2 infection (B) hACE2-APP/PSEN1 mice that resulted in the loss of 10 mice out of 23 suggesting a severe reaction in this strain compared to the hACE-3xTg (C) The Kaplan-Meier curve is plotted for the surviving fraction of hybrid hACE2–AD mice used for molecular MR imaging.

Figure 2B shows the histopathological characterization of 14-month-old hACE2- APP/PSEN1 (Figure 2Bi) and 17-month-old hACE2-3xTg (Figure 2Bii) mouse brains. The development of relevant deposits (amyloid plaque and pTau) in the brain confirms the phenotype of these mice.

The susceptibility of the crossbreeds to SCV2 delta variant infection was tested by infecting the mice by intranasal instillation of 10^3^ pfu. Figure 3 shows the change in the body weights of the infected hybrid mice and the Kaplan-Meier survival graph of each cohort. 3xTg crossbred mice demonstrate 2% to 25% weight loss over 10 days, while APP/PSEN1 crossbreeds demonstrate up to 20% weight loss, with some animals showing measurable weight gains. Four of the 16 hACE2-3xTg mice met euthanasia criteria during the 10-day period (Supplementary Figure 1), while 16 of 27 hACE2- APP/PSEN1 mice met euthanasia criteria over a 40-day period. All hACE2-APP/PSEN1 fatalities occurred in the first 11 days of infection (Supplementary Figure 2). Infection with the Delta variant at this level is therefore a suitable model of severe COVID-19.

Neuroinflammation in hACE2-3xTg mice following SCV2 delta variant infection:

Figure 5A shows the experimental scheme by which infected mice were imaged 10 days post infection. Comparison to the pre-contrast images provides the locations in the brain where the nanoparticle contrast is retained. The circulation half-life of these mMRI probe particles is about 18 hours, and therefore at the 4-day point (∼96 hours) the vast majority of the nanoparticles have been cleared from circulation, and any remaining signal can be reasonably assumed to be due to binding of the particles to the target, analogous to similar studies previously reported^41,42^.

**Figure 4:**
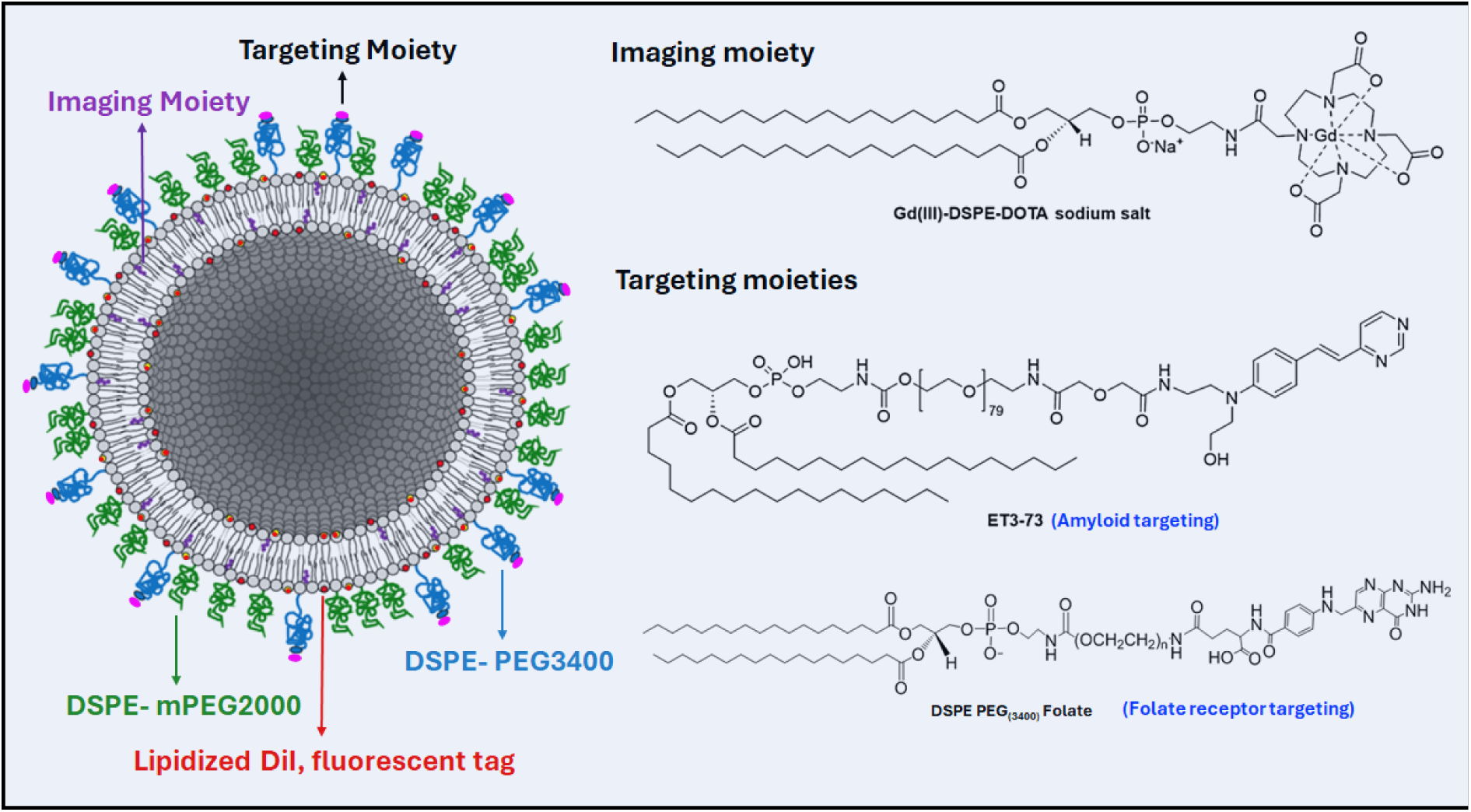
Nanoprobes used for molecular MR imaging. The liposomal bilayer incorporates DSPE-DOTA-Gd (III) for MR contrast readout, lipidized DiC for fluorescence microscopy imaging, DSPE-mPEG2000 to enhance circulation half-life. The control Non-targeted (NT) liposomal formulation contains DSPE-PEG3400 whereas the targeted formulation includes DSPE-PEG3400-ET73 for amyloid beta targeting nanoparticles (AβNP), and (b) DSPE-PEG3400-Folic Acid for the folate nanoparticles (FNP).

**Figure 5:**
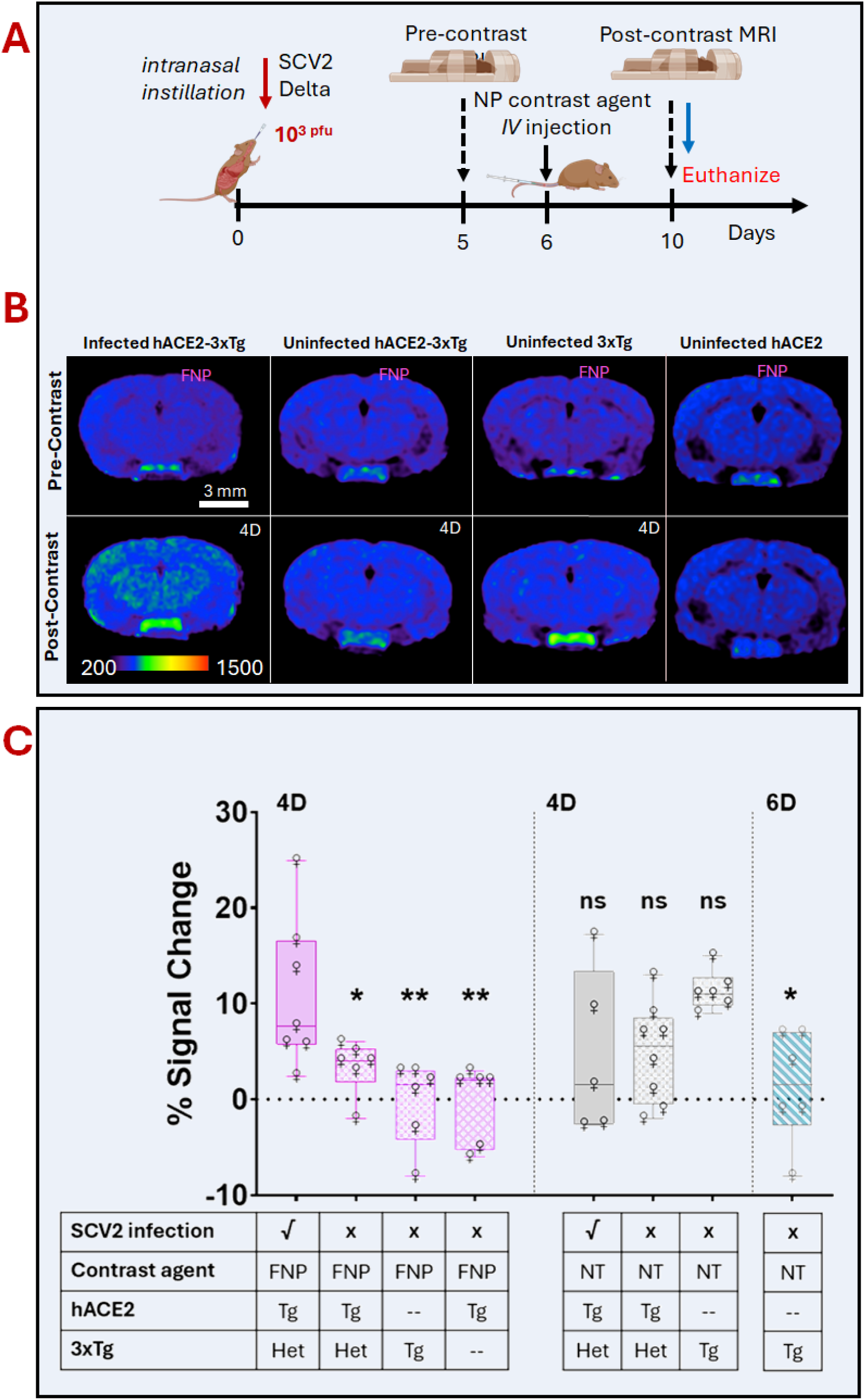
In vivo MR imaging of neuroinflammation in hACE2-3xTg mice (A) Experimental scheme for studies with hACE2-3xTg mouse model. A 10-day short study included infection with SCV2 delta variant at 1x103 PFU. Pre-contrast MRI was performed 5 dpi followed by contrast agent injection 6 dpi and post contrast imaging at 10dpi. Mice were euthanized and their brains harvested for histology. (B) Infected hACE2-3xTg mice demonstrate higher cortical MR signal enhancement relative to uninfected siblings and the parental 3xTg-AD mice at the same age after administration of a folate-targeted nanoparticle. Pre-contrast (PRE) and post-contrast (POST) MR images are shown. Scale bars indicate 3 mm. All pseudo-colored MR images are windowed to the same signal intensity range. (C) Box and whisker plots of MRI signal changes (post-pre injection of probe). The plot also includes the control non-targeted probe (NT) in the infected and uninfected hybrid mice and the parental strain. The post-contrast MR images were also acquired at 6 days post contrast injection for the uninfected hybrid cohort. p-value p <0.05*, p<0.005**, ns-not significant

Figure 5B and supplementary Figure 5 show pre-contrast and post-contrast T1w MRI images for representative infected hACE2-3xTg mice and control groups that include uninfected hACE2-3xTg, 3xTg, k18-hACE2, as well as mice treated with non-targeted probe particles. The non-targeted particles serve as an indicator of vascular leak, as demonstrated in previous work^43^. All images are shown on a uniform color scale, so that visual comparisons can be easily made. Figure 5C shows the quantification of cortical signal enhancement for the test and control groups. It is evident that at the 4-day post- contrast mark, the infected mice exhibit significantly higher signal than their uninfected counterparts. The elevated signal was, however, not statistically significantly different from that observed with non-targeted particles. This is attributed to the slower clearance of the non-targeted particles, as confirmed by the measurement at 6 days post injection, which exhibits a significantly lower signal.

Figure 6 shows histopathological analysis of infected and uninfected hACE2-3xTg mice and 3xTg mice as controls. IBA-1 (a marker of activated microglia), amyloid plaques (stained by 4G8 antibody), pTau (stained by AT8 antibody) and the DiI labeled nanoparticles are all visualized along with cellular nuclei stained by DAPI. Representative images in the cortical and hippocampal areas are shown in Figure 6. The infected mice show elevated levels of microglia activation (IBA-1), amyloid deposits (4G8), and pTau (AT8). There is also a greater retention of targeted nanoparticles in the brains of infected mice, consistent with the increased MRI signal in their brains observed in Figure 5. Data demonstrated increase in NfL levels post-infection is shown in the supplementary Figure 3.

**Figure 6.**
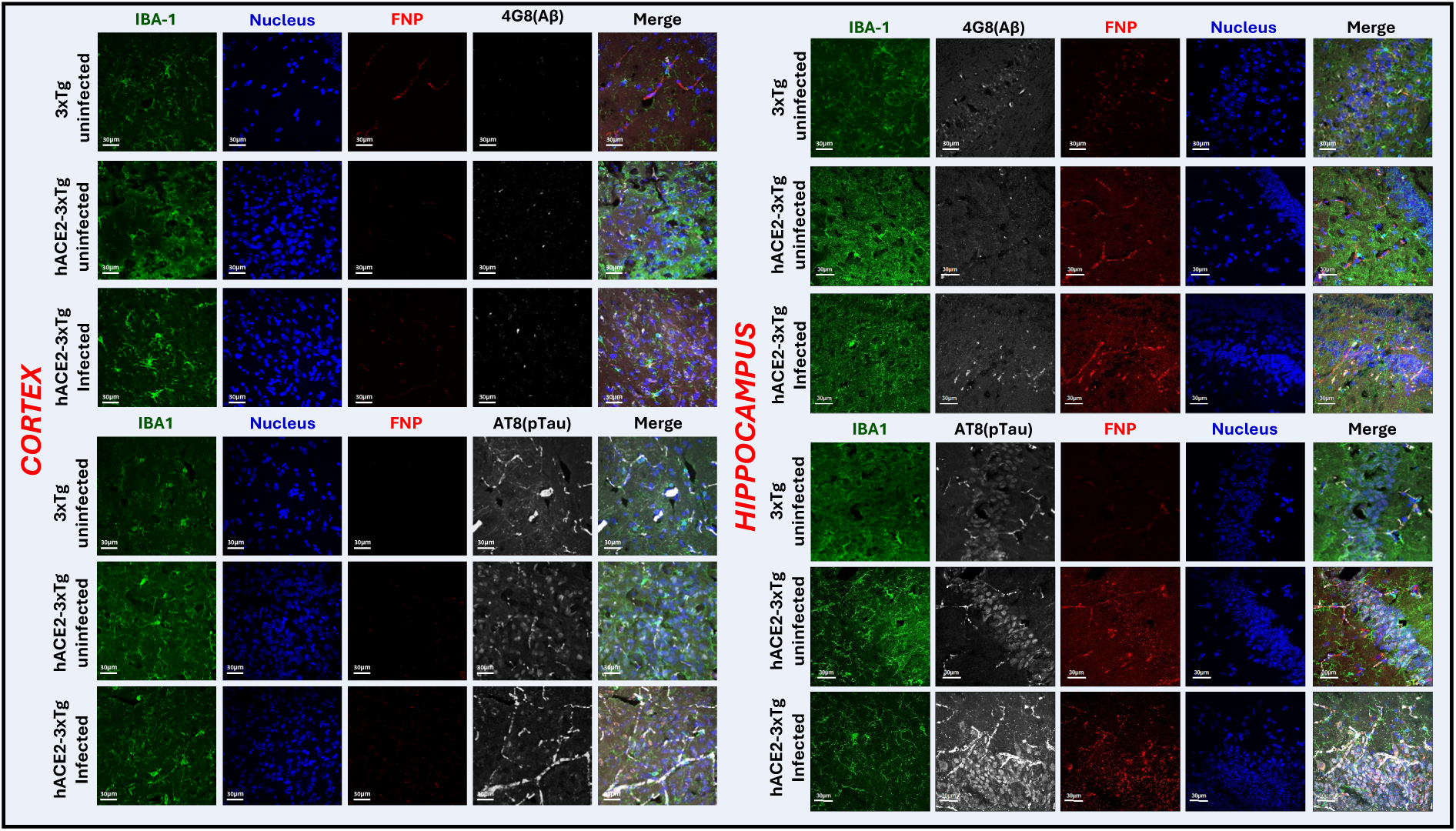
Histological analysis of infected and uninfected hACE2-3xTg and parental 3xTg mice Immunofluorescence with frozen brain sections from the infected hACE2-3xTg, and the uninfected counterparts alongside the parental 3xTg-AD were stained for different antibodies to investigate the changes upon SCV2 infection. AD pathology was traced using amyloid (Biolegend, β amyloid 4G4) and phosphorylated tau (ThermoFisher, AT8). IBA1(Wako Fujifilm, IBA1) stains for activated microglia. Higher expression of Aβ and IBA1 is observed in infected hybrid mice compared to the uninfected counterparts. Representative images from the cortical and hippocampal region show higher signal in the infected group suggesting acceleration of AD phenotype development.

Amyloid deposition post SCV2 infection:

Figure 7 shows the change in amyloid deposition in the brains of young 3-5-month- old hACE2-APP/PS1 mice 30 days after SCV2 infection. At this age, neither the parental nor the uninfected hybrid mice show any amyloid deposition, the process is typically observed at 6-7 months in the APP/PS1 strain. We hypothesized that infection would accelerate amyloid deposition. The infection/imaging scheme is shown in Figure 7A: a pre-contrast scan was collected on day 25 post infection with 1 X 10^3^ pfu SCV2 Delta variant, amyloid targeted nanoparticle contrast (AβNP) injection was on day 26, and post- contrast imaging was on day 30. The resulting pre- and post-contrast images are shown in Figure 7B. Non-targeted nanoparticle contrast agent was injected in control cohorts of mice. These images are shown in Supplementary Figure 6. Figure 7C summarizes the quantified cortical signal difference (post-pre) for each of the experimental cohorts in this experiment. Supplementary Figure 7presents representative regions of interest (ROIs) used for cortical signal segmentation. It is evident that infected hACE2-APP/PSEN1 mice exhibit higher signal difference than any of the uninfected groups, as well as infected mice imaged with non-targeted particles, consistent with the presence of greater amyloid deposits in the brains of infected mice.

**Figure 7:**
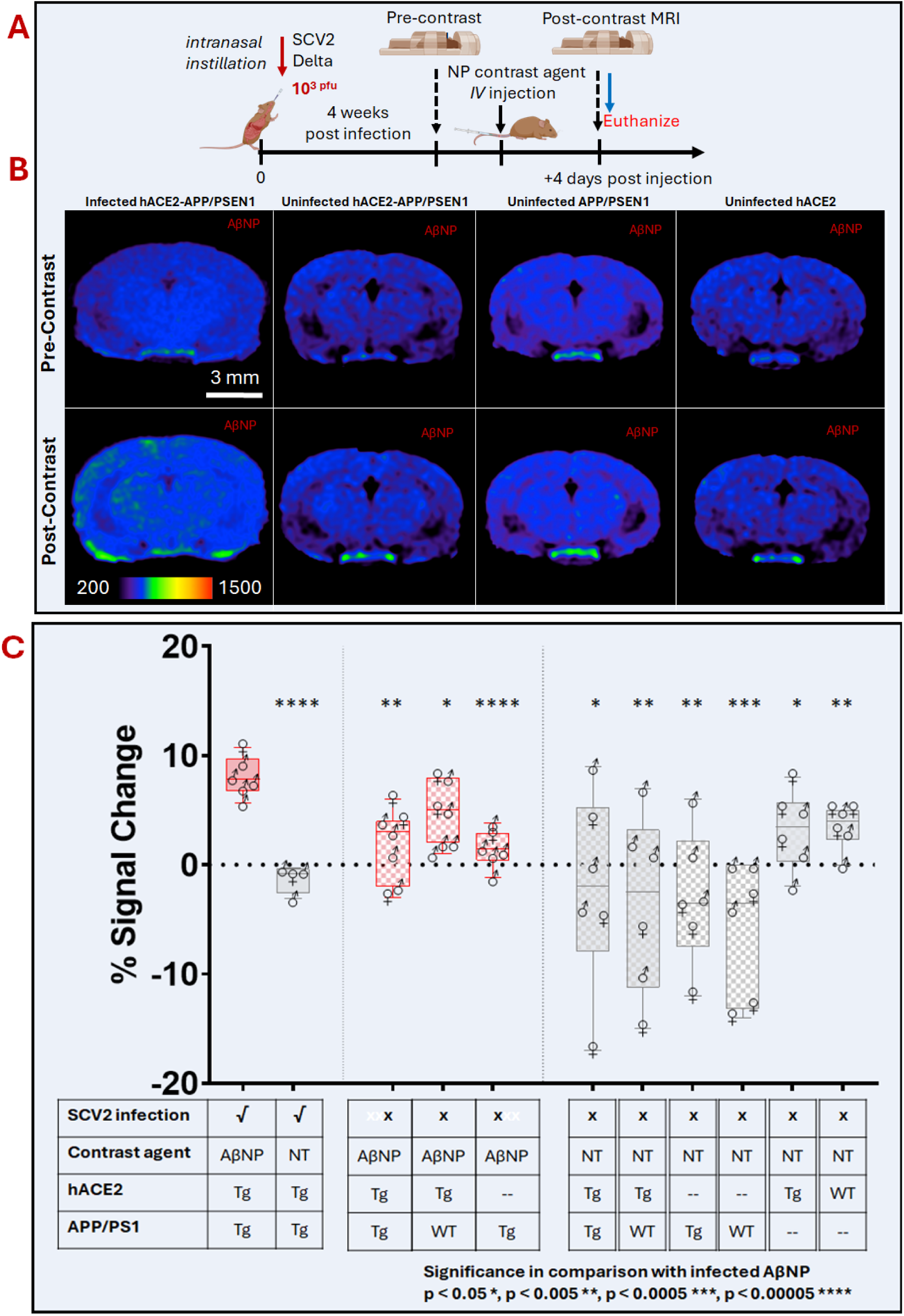
In vivo MR imaging of accelerated amyloid deposition in infected hACE2-APP/PSEN1 mice (A) Scheme of hACE2-APP/PSEN1 mice study. Intranasal SCV2 infection of mice was followed pre-contrast MRI performed 4weeks post infection followed by contrast agent injection and post contrast imaging 4 days post contrast injections. Mice were euthanized and their brains harvested for histology. (B) Contrast enhanced MR images showing signal enhancement in infected transgenic (Tg) k18-hACE/APP-PSEN1 mice. Higher cortical and hippocampal signal relative to uninfected APP-PSEN1 mice after administration of amyloid-targeted liposomal-gadolinium (AβNP) contrast agent. Scale bars indicate 3 mm. All pseudo-colored MR images are windowed to same signal intensity range. (C) Box and whisker plots for the targeted (AβNP) and non-targeted (NT) formulations in the infected and uninfected hybrid mice alongside the parental hACE2 and APP/PSEN1 strain show a significant increase in amyloid in infected mice in comparison with control non-transgenic animals and non-targeted contrast agent. Significance in comparison with infected AβNP p < 0.05 *, p < 0.005 **, p < 0.0005 ***, p < 0.00005 ****

The cortical and hippocampal areas stained for the activated microglia IBA-1, amyloid plaques (4G8), phosphorylated Tau (AT8) show abnormal amyloid plaques in the young 4-month-old, infected animals (Figure 8). Differences are observed in parental and hybrid mice. Higher expression of IBA-1 and Aβ is visualized in fluorescence microscopy. Even though AT8 signal is relatively constant in all the three groups, we see a striking change in the staining pattern of AT8 in uninfected and infected hybrid mice. Similar data for NfL is shown in the supplementary Figure 4.

**Figure 8:**
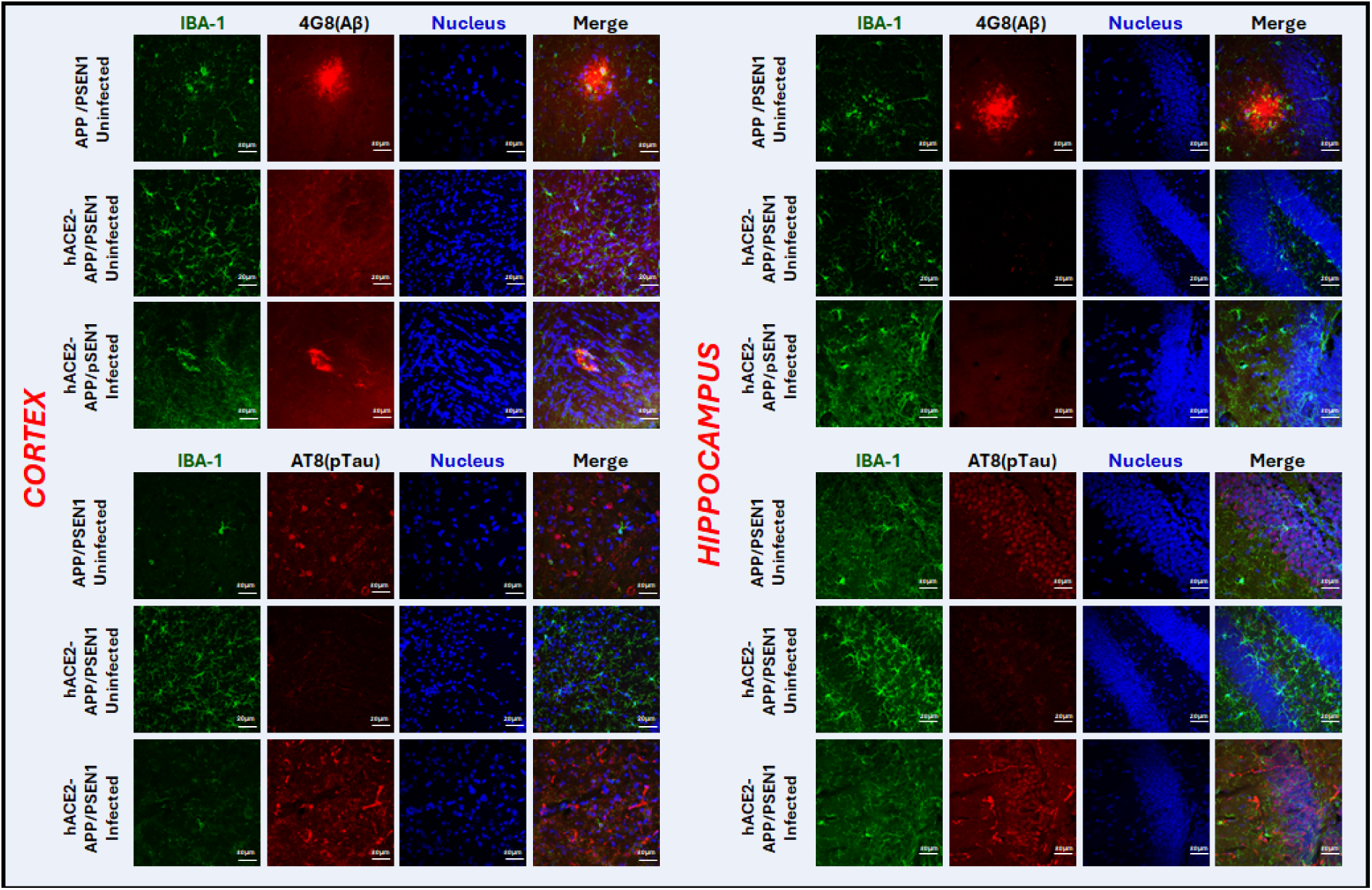
Histological analysis of infected and uninfected hACE2-APP/PSEN1 and the parental APP/PSEN1 Histological analysis of infected and uninfected hACE2-APP/PSEN1 (4 months) and the parental APP/PSEN1 (6 months). Immunofluorescence with frozen brain sections from the infected and uninfected hACE2-APP/PSEN1 hybrid mice and the parental APP/PSEN1 were stained for AD pathology with amyloid (Biolegend, β amyloid 4G4), phosphorylated tau ( ThermoFisher, AT8) antibodies and activated microglia with IBA1(Wako Fujifilm, IBA1) antibody showing accumulation of AD plaques in infected cohort at a younger age (4m) comparable to the parental strain (6m) consistent with the acceleration of AD phenotype development.

## Discussion & Conclusions

The accumulation of amyloid beta^44,45^ and tau^46,47^, alongside chronic inflammation^48,49^, represents three major mechanisms proposed as the contributing factors in the development of Alzheimer’s Disease (AD). The chronic inflammation hypothesis proposes that AD may be triggered by exposure to a pathogen that initiates an inflammatory response, ultimately leading to neurodegeneration^50^. Herpes viruses have previously been studied as possible predisposing factors for AD^51^. The long term neurological components of the post-acute sequelae of SCV2 infection (PASC)^52,53^ highlight the need to explore the biology of SCV2 infections and their potential impact on neuroinflammation and its possible impact on AD development.

Our investigations incorporated a transcriptomic study to determine the overlap of differentially expressed genes involved in AD development and ancestral SCV2 infection. The results provide insights into shared pathways between AD mouse models—tau (P301S), amyloid (APP/PSEN1)—and SCV2 infection in K18-hACE2 mice^25,54,55^. Inflammatory processes, particularly interferon signaling and the complement cascade, emerged as predominant shared pathways. Notably, APP/PSEN1 mice exhibited Type 1 interferon and cell-specific innate immune responses unique to SCV2 infection. Collectively, these findings suggest common inflammatory and cytokine processes between SCV2 infection and AD in these mouse models. These results are consistent with recent results from spatial transcriptomic studies^16^ that also revealed a similar overlap in inflammatory pathways in COVID-19 patients.

To further examine the impact of SCV2 infection on AD pathogenesis, we used SCV2 Delta variant to develop a mouse model capable of both SCV2 infection and AD hallmark formation. The AD mouse models 3xTg-AD^29^ (with both tau and amyloid pathology) and APP/PSEN1^28^ (with only amyloid deposits) were selected and crossed with mice expressing human ACE2, allowing infection with human SCV2 strains. The progeny of these crosses displayed AD pathology phenotypically similar to the parental AD strains. SCV2 infection of the resulting hybrid hACE2-3xTg and hACE2-APP/PSEN1 mice revealed differential susceptibility. The hACE2-APP/PSEN1 mice developed more severe signs of infection, with ∼50% losing over 15% of body weight between five- and nine-days post-infection, making them suitable for severe COVID-19 studies. The surviving mice in both hybrid models were then used to assess SCV2 impact on the development of AD pathology.

We studied 10-month-old hACE2-3xTg mice, prior to the development of amyloid plaques, which typically occur at 14 months in parental strain. In a ten-day study using SCV2 Delta variant, *in vivo* changes were monitored via mMRI using a folate nanoparticle probe (FNP) for imaging of neuroinflammation^56,57^. Folate receptor β has been demonstrated as a marker of inflammation^58^. Low blood folate levels elevate neurotoxic amino acid homocysteine that induces ROS and excitotoxicity which in turn triggers inflammation associated with AD^59,60^. Low folate levels upregulate folate receptors that mediate the intake of cellular folate^61^. We used FNP that bind folate deprived cells that is higher in the pathologic reactive astroglia that shows a higher uptake of the FNP particles^62–64^. Four-days post-FNP injection, infected mice displayed significantly greater contrast enhancement compared to uninfected hybrid and parental strains, indicating neuroinflammation due to SCV2 infection. Comparisons with infected mice using the non- targeted (NT) probe revealed no significant difference, possibly due to variations in circulation times and host interactions, aligning with previous results showing a longer residence time for the NT probe versus folate nanoparticles. We confirmed these findings by conducting mMRI six days post-injection in uninfected hybrid mice, further establishing FNP probe specificity.

In studying the APP/PSEN1 mouse model, which typically shows visible amyloid plaques at 6-7 months, we infected 4-month-old hybrid hACE2-APP/PSEN1 mice to test for accelerated amyloid deposition. mMRI performed using both Aβ-targeted and NT probes four weeks post-SCV2 infection showed significant differences in contrast between infected and uninfected hybrid and parental strains with the Aβ-targeted probe showed higher signal indicating accelerated amyloid deposition at age of 4 months in infected hybrid mice, an age when the parental strain does not show amyloid deposition. The mMRI studies revealed neuroinflammation at 10 days post-infection in the hACE2-3xTg model and, accelerated amyloid deposition at 4 weeks post-infection in the hACE2-APP/PSEN1 model, that were both absent in the corresponding uninfected hybrids. Immunohistochemical validation was done by staining for activated microglia, amyloid pathology, hyperphosphorylated tau, and neurofilament L (associated with neuronal injury). Infected samples, compared with corresponding uninfected and parental sections, showed higher fluorescent signal that corelates with the mMRI findings with the FNP and AβNP probes in the hACE2-3xTg and hACE2-APP/PSEN1 mouse models respectively. Regarding the hACE2-3xTg mouse model, higher signal in the SCV2 infected samples compared to the controls for IBA1 that stains activated microglia confirm neuroinflammation. The hACE2-APP/PSEN1 model showed the presence of amyloid plaques at 4 months of age similar to the initial plaques visible beyond 6 months of age in the parental APP/PSEN1 strain. The higher sensitivity of fluorescence images allowed us to visualize the early plaque formation not visible by mMRI with the AβNP probe and further supports our hypothesis that SCV2 infection led processes results in accelerated deposition of amyloid. Higher intensity fluorescence corresponding to pTau and NfL were also observed in both the mouse models.

In summary, our findings in mouse models of AD and SCV2 infection suggest that SCV2 infection increases neuroinflammation and accelerates amyloid deposition. This study supports further research into the infectious theory of AD pathogenesis and could inform the development of novel therapies and preventive strategies for AD.

## Limitations of studies

This study was conducted at a single time point for both models, which limits the understanding of long-term dynamics in the relationship between SCV2 infection and AD pathology. To uncover a more detailed mechanistic basis for the infection-AD connection, a longitudinal study would be necessary. Additionally, the potential effects of multiple infections and viral strain evolution on AD development remain unexplored. These aspects warrant further investigation to comprehensively assess the infection’s impact on AD progression.

## Author Contributions

Study concept and design: PP, AA. Supervision: AA. Project coordination: PP, AAB. Drafting of original manuscript: PP. Editing -PP, AAB, KG, SER, AKR, AA Experimental - BSL3: AAB, JRR, LJB, ARK, JLC, RB. Nanoparticle Synthesis: RB, PB. mMRI – PP, AAB. Animal Breeding, Necropsy and i.v. injections: RM, MS, RB, XS. Immunohistochemistry: PB, SN, PA, PP. All authors approved the final version of paper.

## Competing Interests

AA, MS, ET hold stock in Alzeca Inc. AA, KG, ET have received consulting fees from Alzeca Inc.

## Acknowledgments

This study was funded by grants from the Barineau, Torian and Levy families. Additional support from the National Institutes of Health (R61/33HD105593) is also gratefully acknowledged. This work was completed using resources from the Texas Children’s Hospital Feigin Pathogen Resource Core and Biosafety Level 3 Laboratory.

## Supplementary Data

**Supplementary Figure 1:**
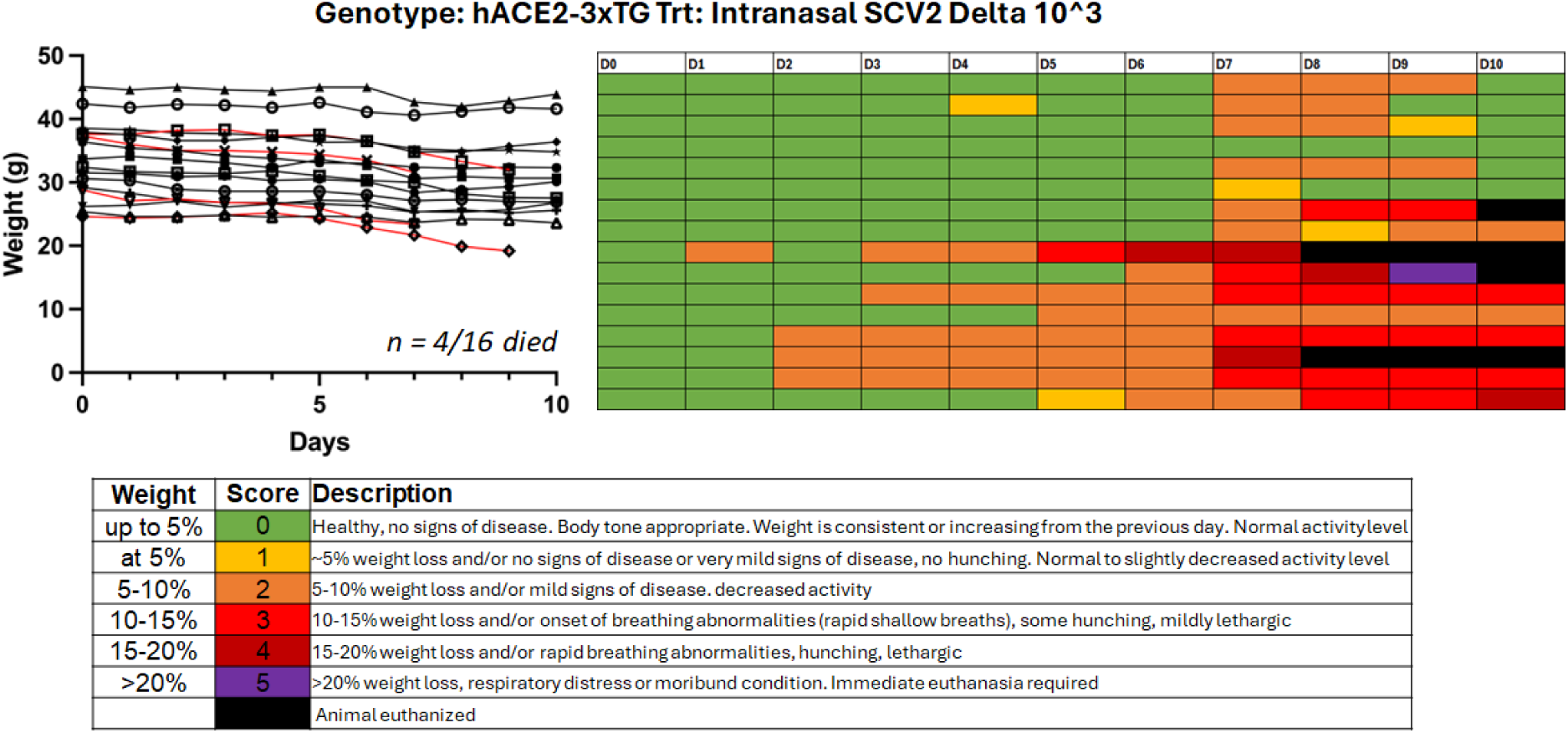
Body conditioning scores and weights of SCV2 infected hACE2-3xTg mice

**Supplementary Figure 2:**
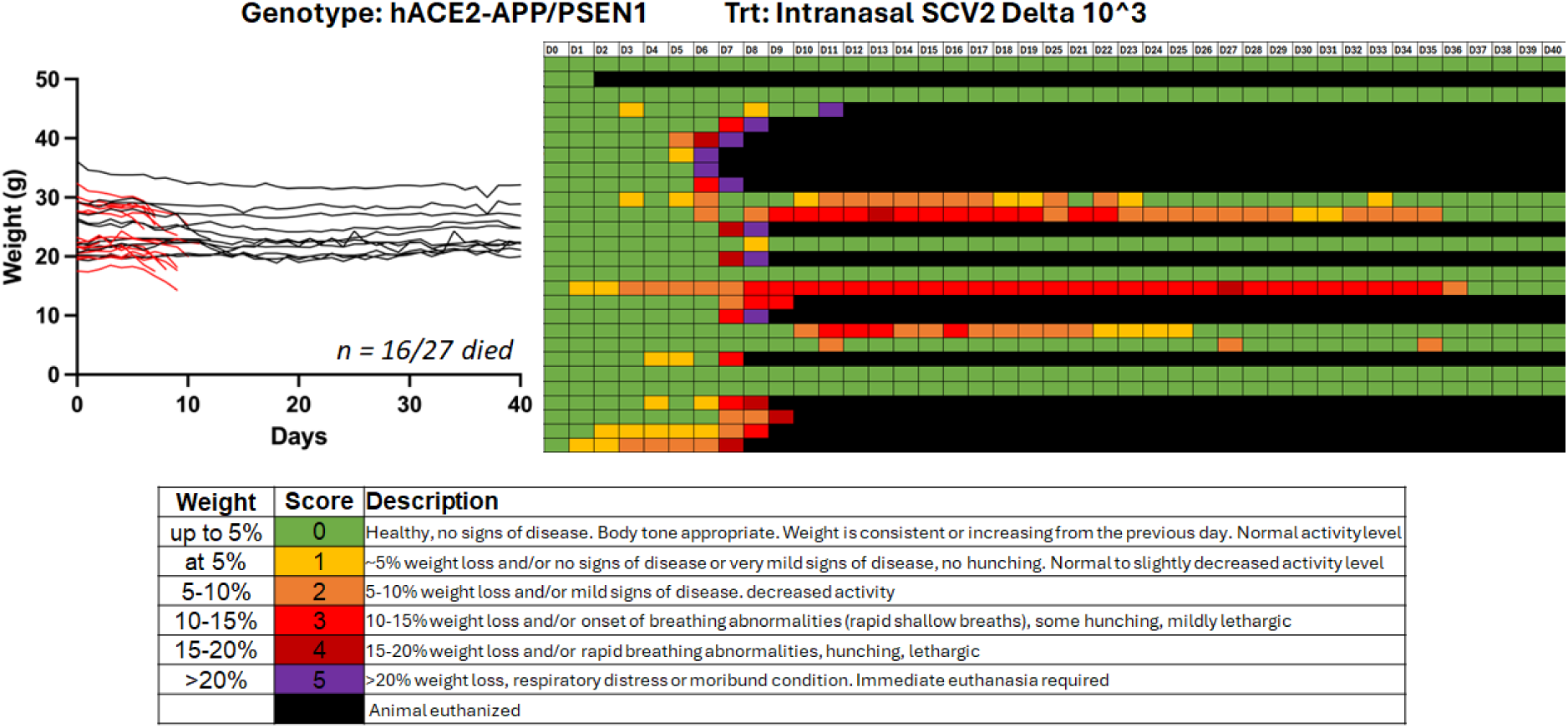
Body conditioning scores and weights of SCV2 infected hACE2-APP/PSEN1 mice

**Supplementary Figure 3:**
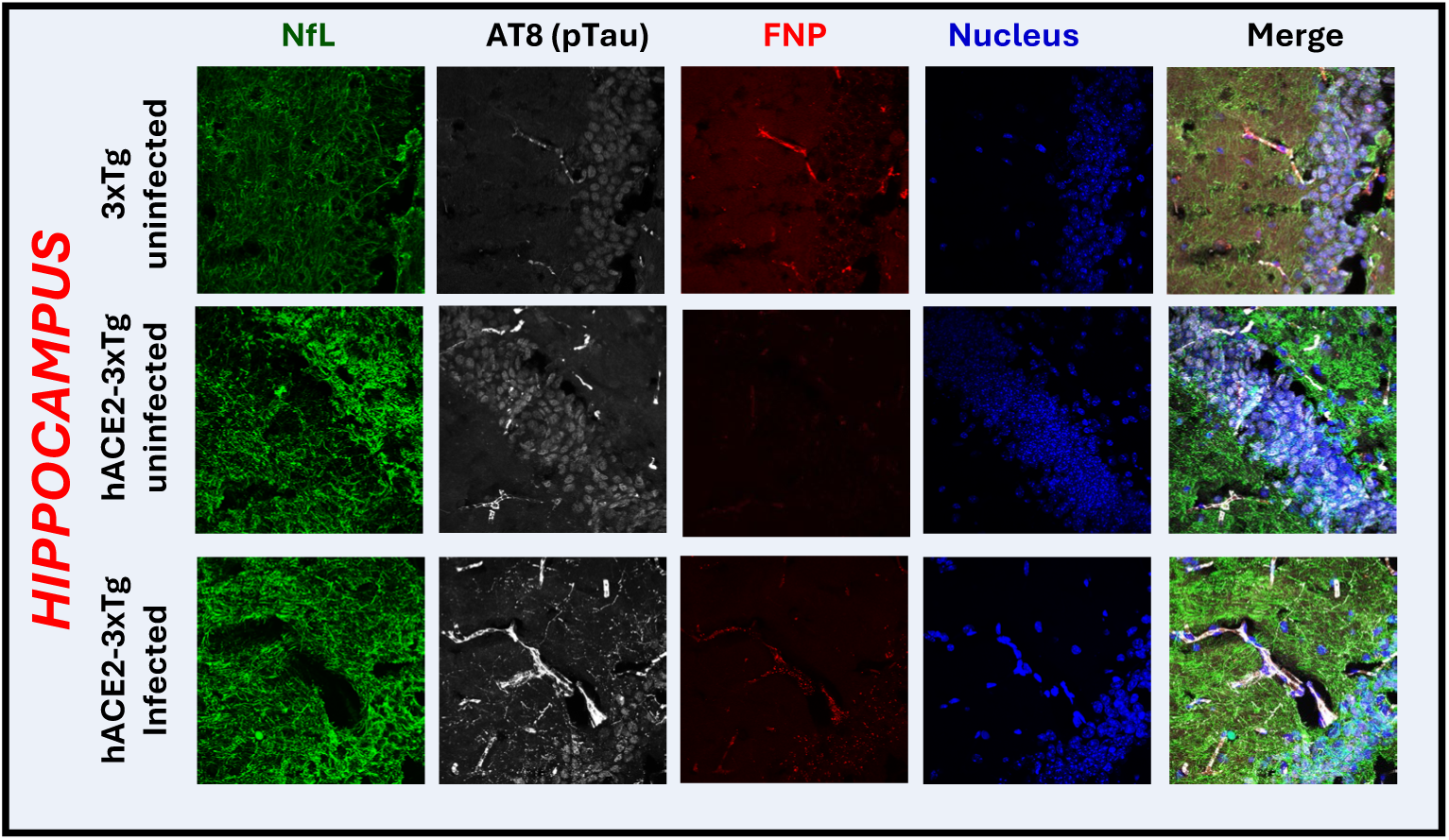
Histological analysis of infected and uninfected hACE2- 3xTg and the parental 3xTg-AD stained with NfL

**Supplementary Figure 4:**
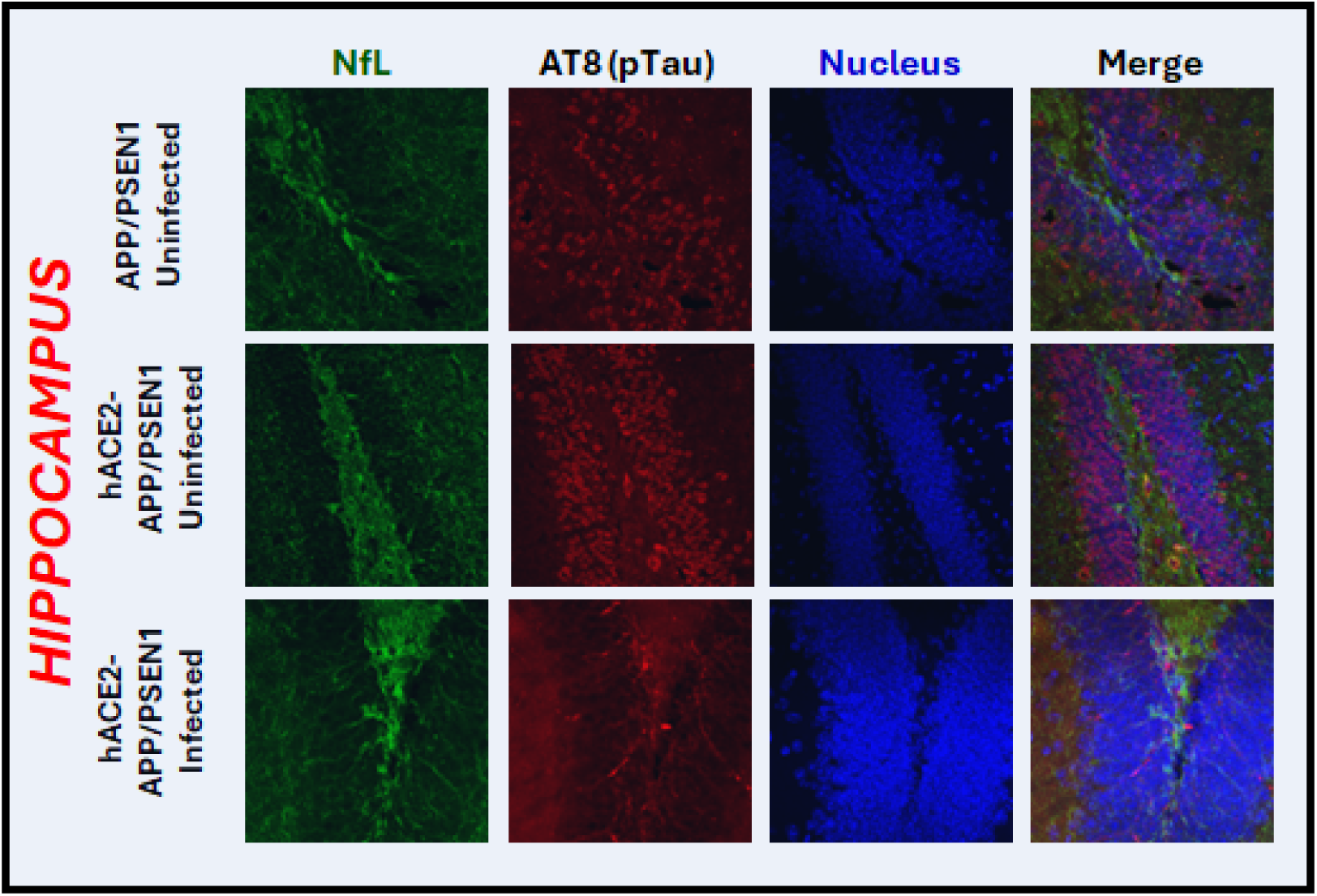
Histological analysis of infected and uninfected hACE2- APP/PSEN1and the parental APP/PSEN1 stained with NfL

**Supplementary Figure 5:**
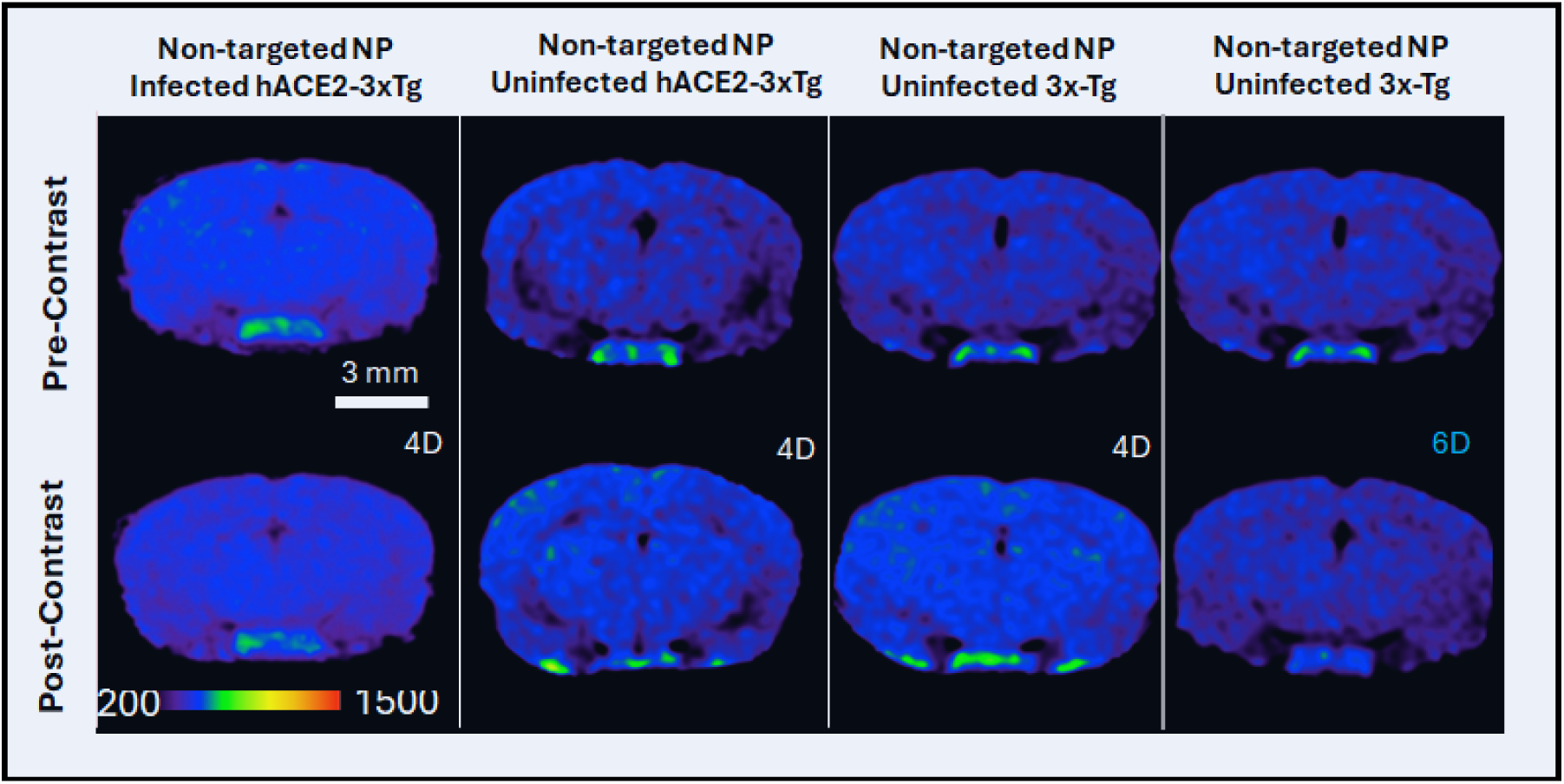
hACE2-3xTg NonTargeted Pre-contrast MRI was performed 5 dpi followed by contrast agent injection 6 dpi and post contrast imaging at 10dpi. No cortical signal enhancement in infected hACE2-3xTg mice relative to uninfected siblings after administration of a non-targeted nanoparticle. Pre-contrast (PRE) and post-contrast (POST) T1W-MR images in 3xTg mice administered folate-targeted preps and hACE2-3xTg mice administered folate-targeted preps are shown. Scale bars indicate 3 mm. All pseudo-colored MR images are windowed to same signal intensity range.

**Supplementary Figure 6:**
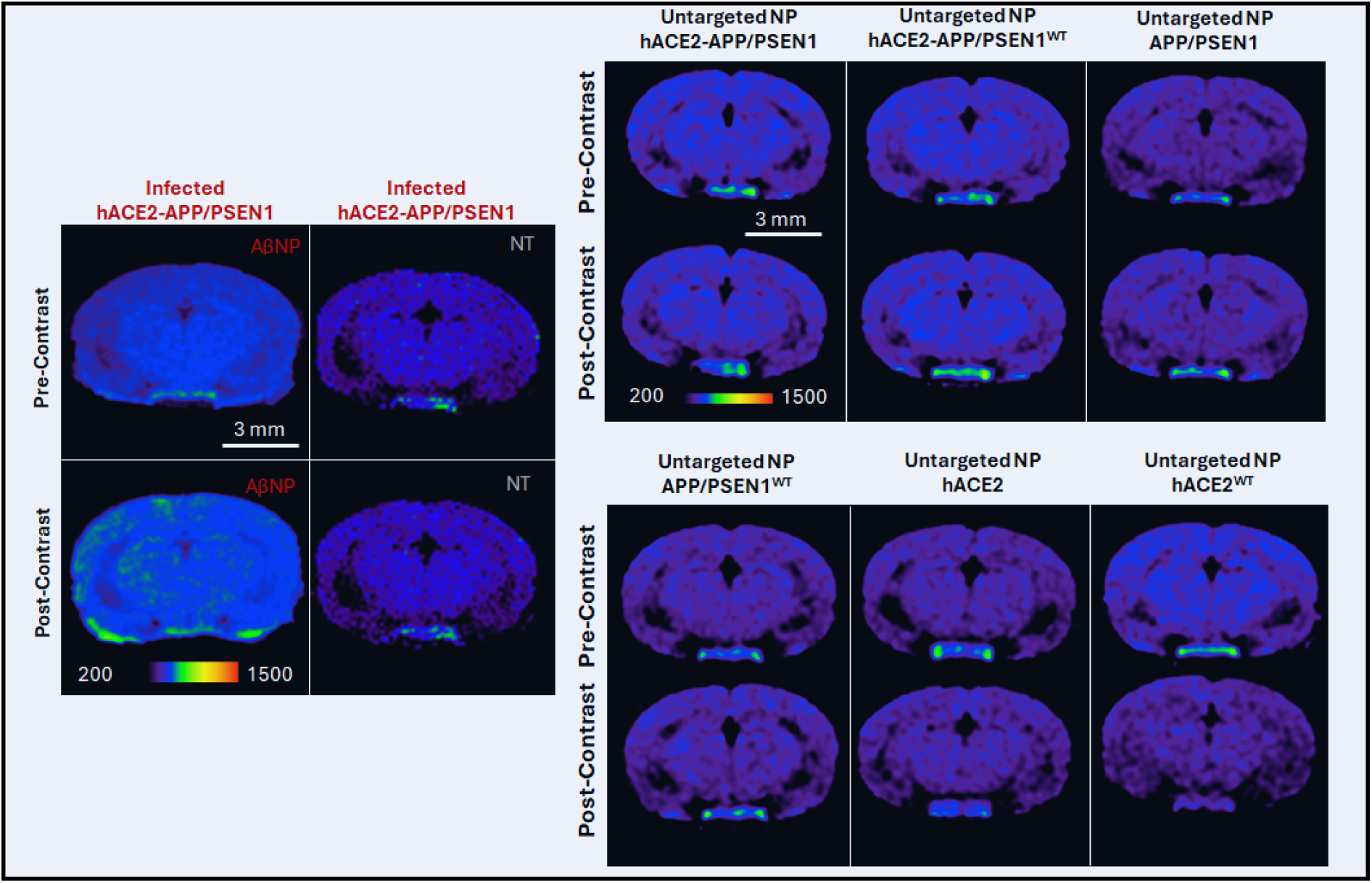
hACE2-APP/PSEN1 NonTargeted Intranasal SCV2 infection of mice was followed by pre-contrast MRI performed 4weeks post infection, followed by contrast agent injection and post contrast imaging at 1 month post infection. Mice were euthanized and their brains harvested for histology. No signal enhancement in infected relative to uninfected APP-PSEN1 mice after administration of non-targeted liposomal-gadolinium contrast agent. Scale bars indicate 3 mm. All pseudo- colored MR images are windowed to same signal intensity range.

**Supplementary Figure 7:**
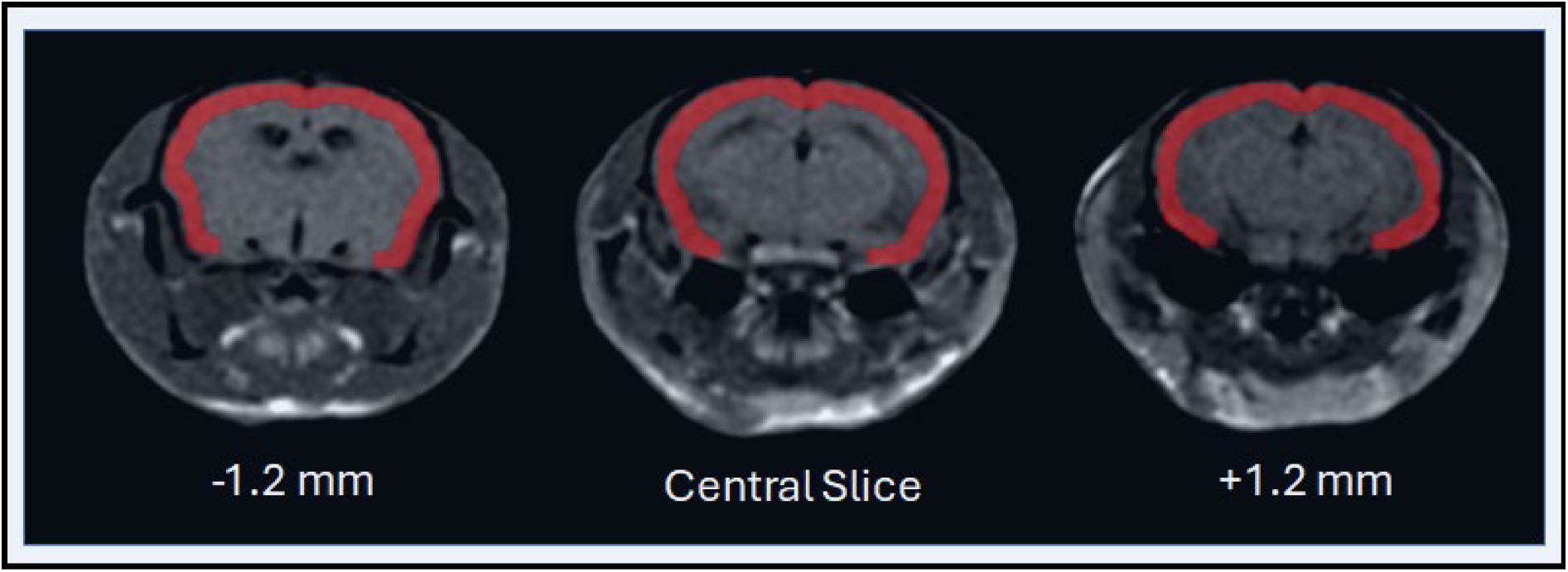
Segmentation of cortical signal using OsiriX (version 5.8.5, 64-bit) List of common DEG’s in K18 SCV2 and P301S mouse model in Figure 1 A (iii) Heat map showing the intersection of differentially expressed gene sets (corrected p<0.05) between infected and uninfected mice, and between P301S transgenic and wild type mice. (iii) SCV2(1) and P301S(2)

